# FOCAL3D: A 3-dimensional clustering package for single-molecule localization microscopy

**DOI:** 10.1101/777722

**Authors:** D. Nino, D. Djayakarsana, J. N. Milstein

## Abstract

Single-molecule localization microscopy (SMLM) yields an image resolution 1-2 orders of magnitude below that of conventional light microscopy, resolving fine details on intracellular structure and macromolecular organization. The massive pointillistic data sets generated by SMLM require the development of new and highly efficient quantification tools. Density based clustering algorithms, such as DBSCAN, can provide spatial statistics on protein/nucleic acid aggregation or dispersion while explicitly identifying macromolecular clusters. The performance of DBSCAN, however, is typically dependent upon an arbitrary, or at least highly subjective, parametric tuning of the algorithm. Moreover, DBSCAN can be computationally expensive, which makes it arduous to evaluate on large image stacks. This is all the more important in 3-dimensions where there exist limited alternatives for quantifying clustering in SMLM data, and where a 2-dimensional analysis of true 3-dimensional data may give rise to image artefacts. We have developed an open-source software package in Python for both identifying and quantifying spatial clustering in 3-dimensional SMLM datasets. FOCAL3D is an extension of our previously developed, 2-dimensional, grid based clustering algorithm FOCAL. FOCAL3D provides a highly efficient way to spatially cluster SMLM datasets, scaling linearly with the number of localizations, and the algorithmic parameters may be systematically optimized so that the resulting analysis is insensitive to variation over a range of parameter choices. We initially validate the performance and parametric insensitivity of FOCAL3D on simulated datasets, then apply the algorithm to 3-dimensional, astigmatic dSTORM images of the nuclear pore complex in human osteosarcoma cells.

The data and software package are available at: http://www.utm.utoronto.ca/milsteinlab/software/

## 1 Introduction

Single-molecule localization microscopy (SMLM) techniques such as photo-activated localization microscopy (PALM) ([1]), super-resolution optical fluctuation imaging (SOFI) ([2]), direct stochastic optical reconstruction microscopy (dSTORM) ([3]), and so on, have revolutionized the way we see the biological world. These techniques can resolve cellular features 1-2 orders of magnitude smaller than what is achievable by conventional, diffraction limited light microscopy. Typical lateral resolutions are approximately 10-20 nm with slightly poorer depth resolution, although recent advances suggest this may be further reduced by another factor of 10 ([4, 5]). Considering that nucleic acids such as DNA or RNA are 2 nm wide and that a typical protein is on the order of 4 − 6 nm in diameter, light microscopy will soon encroach upon the domain of electron microscopy, enabling us to quantify cellular organization and physiology down to the scale of single macromolecules.

Images acquired by conventional fluorescence microscopy are diffraction limited, spatial maps of fluorophore intensity while SMLM yields a pointillistic set of approximate single-molecule coordinates (i.e., localizations). There are various approaches to generating image reconstructions from SMLM datasets that appear to produce images analogous to conventional microscopy, albeit with an enhanced resolution. For instance, one can render a Gaussian with a width of the localization precision at each localization coordinate, but the resulting image is not actually an intensity map, rather it represents the probability density of finding a single-molecule at a given location. Regardless, image processing techniques used to analyze conventional microscopy images are often applied to SMLM image reconstructions. A more quantitative approach, however, is to work directly with the table of localizations.

Various techniques exist to analyze statistical properties of pointillistic data sets and have already been applied to analyze SMLM data, such as pair-correlation analysis ([6]) or the Ripley’s K-function ([7]). These ensemble measures are able to quantify statistical properties of the localization data such as the degree of clustering or the average size of a cluster, but do not directly identify individual clusters within a dataset. Density based spatial clustering algorithms, on the other hand, attempt to directly assign groups of localizations to a single cluster and can provide information both on the distribution of cluster size and shape as well as quantify intracellular, spatial organization.

Density based spatial clustering with noise (DBSCAN) is perhaps the most canonical of these methods ([8]). How DBSCAN performs is determined by two user-defined parameters: a length-scale *ϵ* specifying the neighbourhood in which to define a local density, and a density threshold *minPts* that determines if a point is part of a cluster or not. Selection of *ϵ* and *minPts* is often a subjective process whereby the user tunes the parameters until DBSCAN does a reasonable job identifying clusters in the data. Tuning of the parameters requires DBSCAN to be run multiple times on each dataset, a process that can be arduous for large SMLM datasets *(n*_*l*_ > 1 × 10^6^ localizations) since DBSCAN scales on average like n_*l*_ log n_*l*_ ([8]). To address these issues, a variety of ‘parameter free’ clustering algorithms have recently appeared in the literature that are specifically designed for SMLM.

Originally developed for 2-dimensional datasets, ([9]) employ Bayesian analysis to optimize cluster selection, whereas ([10]) take a geometrical approach that makes use of Voronois tessellation. While 2-dimensional algorithms are useful for analyzing clustering in a plane, say within the cellular membrane, many biological systems require a true 3-dimensional analysis and simply clustering a projection of the data will often lead to false artefacts. In response, both of these methods have recently been extended to handle 3-dimensional datasets ([11, 12]). The runtime of these algorithms, however, is quite long, requiring several hours to evaluate a single analysis on a reasonably large set of 3-dimensional SMLM data.

Here we present FOCAL3D, a 3-dimensional implementation of our previously released Fast Optimized Clustering Algorithm for Localization Microscopy (FOCAL) ([13]). FOCAL3D is a density based approach similar to DBSCAN, but performed upon an appropriately discretized spatial grid. Under most conditions, this discretization greatly speeds up the algorithm, which scales linearly with the number of localizations *n*_*l*_, enabling the user to rapidly optimize and identify clusters in SMLM data, facilitating the throughput of image analysis. While FOCAL3D is written in Python, both the source code and a Windows executable file are made available, providing users with a simple graphical user interface (GUI) so it can easily be implemented by those with no computer programming experience.

An overview of this manuscript is as follows. In Section 2.1 we provide background on the FOCAL3D algorithm and, in Section 2.2, discuss how to optimize the algorithmic parameters. In Section 3.1 we discuss our SMLM simulations and, in Section 3.2, we evaluate the performance of FOCAL3D in clustering the simulated data at both moderate and high noise conditions. Then in Section 3.3 we assess the clustering of FOCAL3D applied to 3D SMLM images of the nuclear pore complex (NPC). We conclude with a discussion of our results in Section 4.

## 2 Methods

### 2.1 FOCAL3D Algorithm

FOCAL3D follows a similar algorithm to the 2-dimensional implementation elaborated in ([13]). Starting from a localization table, each localization coordinate is first assigned to a discrete spatial bin (voxel) of volume Δ^3^. This creates a discretized localization density map. To enhance contrast between the noise and target (i.e., cluster) points, rather than use the raw density map, we assemble an enhanced density map by replacing the value at each voxel by a sum over all neighbouring voxels (i.e., a 3×3 grid in 2D or a 3×3×3 cubic grid in 3D centred on the voxel under consideration). This contrast enhancement is performed only for bins that contain at least one localization.

Candidate clusters are selected by first identifying voxels that have a value above a density threshold *minL*. These voxels are then labeled as core voxels. Voxels adjacent to any one of the faces of a core voxel, but that did not satisfy the density threshold, are classified as border voxels. These core and border voxels comprise the candidate clusters. The final step then is to narrow down the field of candidates by imposing a threshold *minC* on cluster size, where *minC* is simply the minimum number of connected voxels containing core and border voxels that will be accepted as a cluster.

### 2.2 Parameter Selection for FOCAL3D

The overall performance of FOCAL3D is dependent upon an appropriate choice of three user-defined parameters: Δ, *minL* and *minC*. The density threshold *minL* and cluster size threshold *minC* are the discrete analogs of *minPts* and *ϵ* in DBSCAN, respectively (see Table 1). By discretizing space in FOCAL3D, we greatly enhance the speed at which we can evaluate the local density and, therefore, the full clustering of a dataset. This speed up comes at a cost, namely, the introduction of an additional parameter Δ specifying the grid size. Δ was chosen in ([13]) based on the localization precision of the SMLM data, which reflected an uncertainty inherent in the imaging method. Here we relax this criterion and show that by properly tuning Δ, the performance of FOCAL3D displays regions of insensitivity to the choice of *minC*, and that within these regions one generally obtains a best estimate of the ideal clustering. A further discussion on some practical aspects of implementing this optimization is provided in (Supplementary Section 1).

**Table 1:** A guide to the parameters used throughout the manuscript. Units are indicated in parentheses (with ℓ indicating the units of length).

|  | Density Threshold<br>(# of localizations) | Local Volume | Grid Spacing |
| --- | --- | --- | --- |
| DBSCAN | *minPts* | $(4\pi/3)\epsilon^3 (\ell^3)$ | - |
| FOCAL3D | *minL* | *minC* (voxels) | $\Delta (\ell)$ |

#### 2.2.1 Selecting the Density Threshold (*minL*)

For each choice of grid size Δ and cluster size threshold *minC*, it’s necessary to set an optimal density threshold *minL*\*. This is achieved by first randomly scattering the table of localizations within the same volume as the original data. We then repeatedly run FOCAL3D on this random dataset over a range of values of *minL*, keeping Δ and *minC* constant. The optimal density threshold *minL*\* is chosen as the lowest threshold value where FOCAL3D returns zero clusters for the randomly scattered data (see Fig. 1 inset).

**Figure 1:**
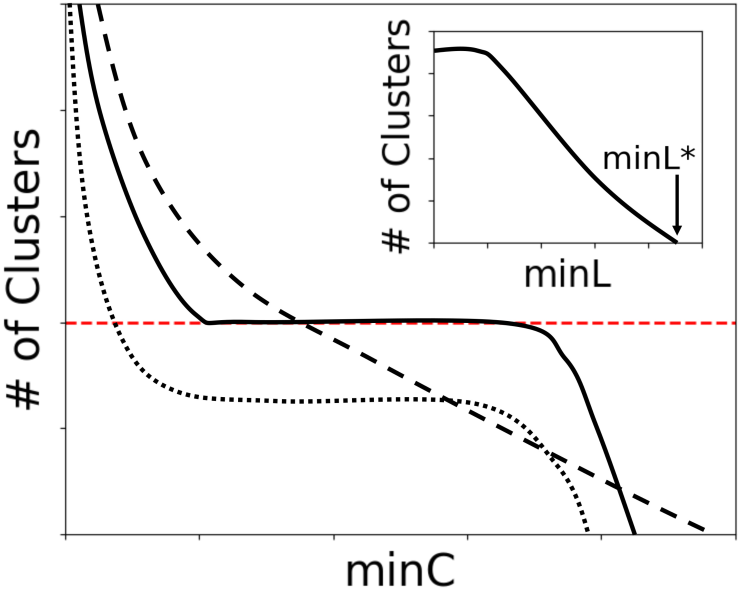
A graphical illustration of the typical behaviour seen in plots of the number of clusters vs. *minC*. Close to an ideal grid size Δ*, regions of insensitivity to *minC* appear (solid line). For Δ ≪ Δ*, a steep dependence on *minC* is often observed (dashed line), while for Δ ≫ Δ* a plateau in *minC* may persist, but generally yields an underestimation in the number of clusters (dotted line). The target number is indicated by the horizontal line (red dashed). Note, *minL* is tuned at each point along the curves to differentiate from a random background (inset).

#### 2.2.2 Tuning the Grid Size (Δ) and Cluster Size Threshold (*minC*)

We next concurrently optimize the grid size Δ and the cluster size threshold *minC*. Toward this end, a series of plots at varying Δ are generated depicting the number of detected clusters vs. *minC* (now, obviously, on the original localization table). Figure 1 illustrates the general trend of these plots where, for a range of grid sizes about an optimal value Δ*, a relatively flat plateau appears. At this choice of grid size, the clustering displays the largest region of insensitivity to *minC*, and along the plateau FOCAL3D does its best job of clustering the localization data and predicting the number of clusters. The optimal grid size tends to fall below the cluster radius Δ* < *R*_*C*_, with too small a choice leading to an overly sensitive dependence on *minC*, and too large a choice resulting in a significant number of missed clusters.

Note that it is not necessary to know the mean cluster size *R*_*C*_ *a priori*, but a good estimate can narrow the range of values for Δ that need to be searched, significantly decreasing the computational time required to optimize the algorithm. For instance, statistical models such as the Ripley’s K-function can yield a reasonable starting point for the search and can be rapidly calculated on localization data. As we will see, there is some flexibility in the choice of grid size that will result in an accurate performance by the clustering algorithm.

Interestingly enough, this parameter selection routine can be translated to DBSCAN. In this case, *minPts* is found from analyzing a random distribution of the original localizations and set once the algorithm no longer detects clusters. Then, since there is no equivalent to the grid size Δ in DBSCAN, a single plot of the number of clusters detected vs. *ϵ*, optimizing *minPts* at each point, is generated and *ϵ* is set within the flat region of the curve.

## 3 Results

### 3.1 Clustering Simulations

We performed a series of analyses on simulated SMLM clustering data to quantitatively assess the performance of FOCAL3D. Here we considered the case of spherically symmetric clusters of mean radii 40±8 nm, 60±12 nm and 80±16 nm, within a cubic volume of 125 µm^3^. As this method is not designed to deal with spatial overlap, the centroids of the clusters were separated by at least 330 nm for the 80 nm clusters, 310 nm for the 60 nm clusters, and 290 nm for the 40 nm clusters. Each cluster consisted of a Poisson-distributed number of ‘dyes’ randomly placed, with uniform probability, throughout the cluster volume (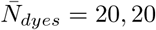 and 10 for the 80, 60, and 40 nm clusters, respectively). To account for blinking, each dye then yielded an exponentially distributed number of localizations 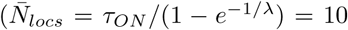 (Supplementary Section 2.1) for all cluster sizes, where *τ*_*ON*_ is the average ON time of a blink and *λ* is the characteristic number of blinks of a fluorophore). Each blink was in turn scattered about the dye centre with a Gaussian distribution whose width was sampled from a distribution of localization precisions (Supplementary Section 2.2). We assumed that the spread in localization precision was slightly poorer in the axial direction (*δ*_*z*_ = 20 nm) than in the lateral plane (*δ*_*x,y*_ = 10 nm) reflecting the reduced axial resolution of SMLM.

Throughout this manuscript, we will quantify the performance of FOCAL3D in clustering the simulated data both in terms of precision and recall, which are standard metrics of the performance of a clustering algorithm. Precision is defined as the fraction of localizations identified as being part of a cluster that were correctly identified (*P* ≡ TruePositives/(TruePositives + FalsePositives)), while recall measures the fraction of clustered localizations identified by the algorithm (*R* ≡ TruePositives/(TruePositives + FalseNegatives)). By taking the weighted average of the precision and recall, these two metrics can be combined into a single measure called the *F*_1_ score, where *F*_1_ = 2*PR*/(*P* + *R*).

Another useful metric is the silhouette score *S*_*C*_, which quantifies the degree of overlap in the detected clusters. Formally, this is defined as 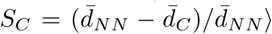, where 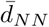 is the mean nearest-neighbour cluster distance and 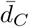 is the mean intra-cluster distance for a sample. *S*_*C*_ will range from 0 to 1 with a higher score indicating more definition and spatial separation of the clusters.

### 3.2 Performance Clustering Noisy Data

We define a measure of the noise in our simulations as follows:

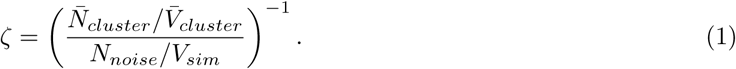

The quantity in parentheses is the level of signal-to-noise, given here by the ratio of the mean density of clustered localizations to the density of noise. 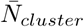 is the mean number of localizations within a cluster, 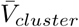 is the mean volume of a cluster, *N*_*noise*_ is the total number of localizations uniformly added as noise to the simulation, and *V*_*sim*_ is the total simulation volume. A reasonable range of values for *ζ* to consider runs from *ζ* = 0, where there’s no noise, to *ζ* = 1 where the density of the noise is equivalent to the density of the clusters.

#### 3.2.1 Optimization at Moderate Noise Levels

Figure 2 displays the number of clusters detected by FOCAL3D as a function of *minC* for a noise level of *ζ* = 0.01 and mean cluster radii of 80 nm, 60 nm and 40 nm. For 80 nm clusters, we now clearly see the behaviour illustrated in Fig. 1 empirically. In this case, for a grid size of half the mean cluster radius (Δ = 40 nm), the number of clusters found by FOCAL3D displays an extended plateau in the neighbourhood of the target number, providing a range of values for *minC* that yield optimal results. However, as *minC* is decreased below this optimal range, the algorithm detects an increasing number of small clusters, which leads to a severe overestimation of the number of target clusters. Likewise, as *minC* is elevated beyond the plateau, FOCAL3D begins rejecting ever larger clusters and the number of clusters predicted by the algorithm steadily decreases to zero.

**Figure 2:**
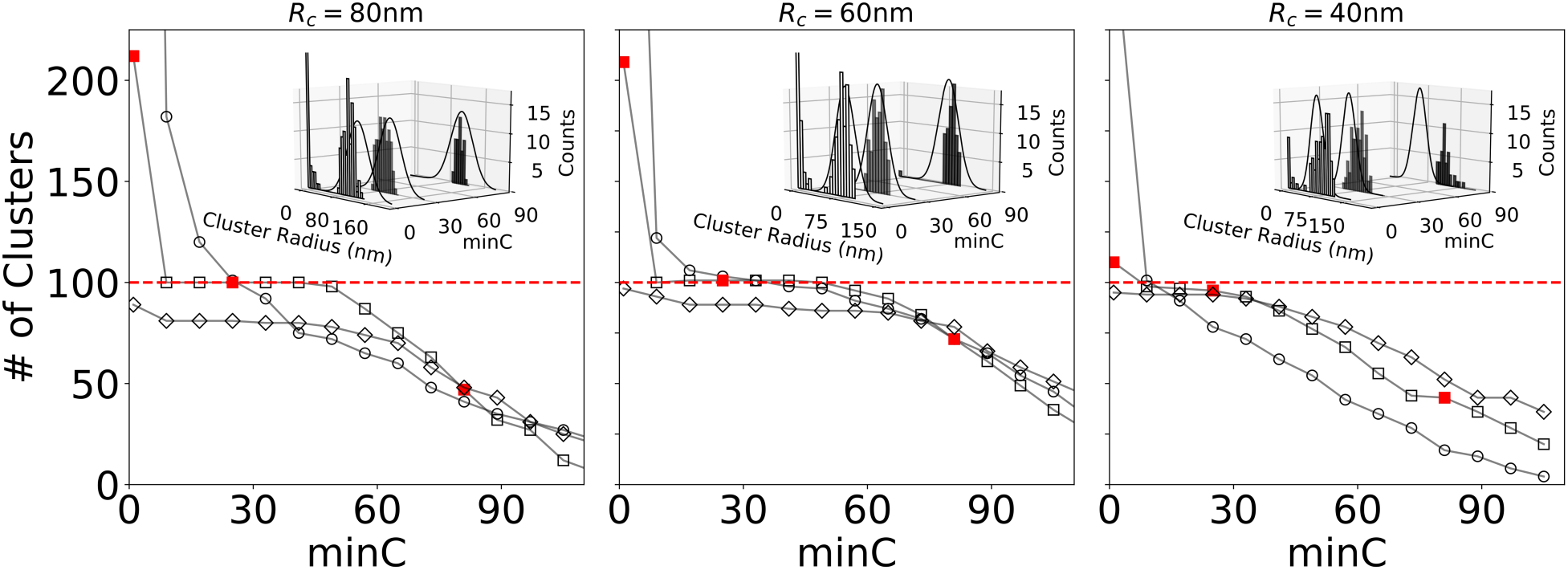
FOCAL3D performance for clusters of mean radius *R*_*c*_ = 80 nm, 60 nm and 40 nm at a noise level of *ζ* = 0.01. At each radius, the number of clusters as a function of *minC* is displayed for a range of grid sizes (*R*_*c*_ = 80 nm: Δ = 20 nm (○), 40 nm (□), 90 nm (◊); *R*_*c*_ = 60 nm: Δ = 20 nm (○), 30 nm (□), 70 nm (◊); *R*_*c*_ = 40 nm: Δ = 20 nm (○), 40 nm (□), 60 nm (◊)). Insets: Distribution in cluster radii estimated from convex hull determined at three points along *minC* (indicated by solid red squares). For comparison, we also display the ground truth distribution in cluster radii, shown by the solid black curves (Supplementary Fig. S5).

If we then adjust the grid size so that too fine of a grid is chosen (Δ = 20 nm), the plateau disappears and the performance of the algorithm becomes strongly dependent on the choice of *minC* (Fig. 2). While the curve does intersect with the target value, without knowing the number of clusters *a priori*, it would be challenging to select the appropriate value of *minC*. Likewise, if the grid is chosen to be too coarse (Δ = 90 nm), an insensitivity to the choice of *minC* may appear, but the algorithm tends to miss a significant amount of clusters within this region of parameter space. We should note that, to obtain the plots in Fig. 2, we scanned through a range of grid sizes Δ (in 10 nm steps) and selected for our optimized choice the one that displayed the most extended plateau at the target. For clarity, not all grid sizes are shown (Supplementary Fig. S6).

We also find that the calculated F1 scores, displayed in Fig. 3, are consistently the highest along the plateau at the optimized grid size (Δ* = 40 nm), and are significantly poorer within this same region for both the coarse (Δ = 90 nm) and fine (Δ = 20 nm) grids. Furthermore, the F1 scores remain relatively constant along the plateau.

**Figure 3:**
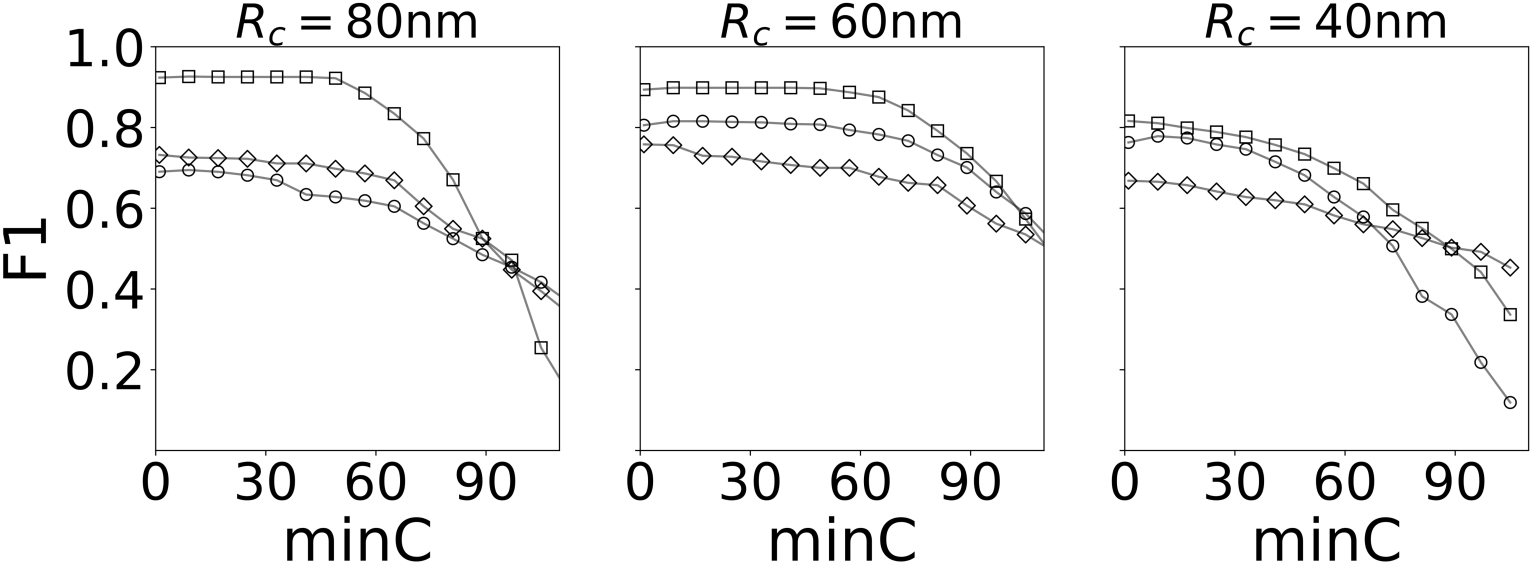
Clustering performance as indicated by the F1 scores (corresponding to the analysis in Fig. 2). The plots show that the algorithm is most correct at identifying clusters within the plateau region (compare to Fig. 2) near a grid spacing Δ*. Symbols the same as indicated in Fig. 2.

Similar behaviour is observed for smaller clusters, with the results for 60 nm and 40 nm clusters shown here (Fig. 2). We note that as the mean cluster size is decreased, the region of insensitivity of the algorithm to *minC* diminishes. For 40 nm clusters, and an appropriate choice of grid size, a plateau is still visible, if just, in a plot of the number of clusters vs. *minC*. We also note that, due to the reduced localization precision along the z-axis, the 40 nm clusters are significantly more elongated than the 80 nm clusters. This tends to shift the choice of ideal grid size to larger values as seen in the figure. Moreover, in both cases the F1 scores are highest, and relatively constant, along the plateau region of the Δ* curve (Fig. 3).

#### 3.2.2 DBSCAN at Moderate Noise Levels

We next consider how DBSCAN performs on the same simulated data sets with an analogous optimization of input parameters (a comparison of run-time scaling is given in (Supplementary Fig. S7)). In Fig. 4, for the case of 80 nm clusters, we plot the number of clusters detected by DBSCAN as a function of *ϵ*, which corresponds to *minC* in FOCAL3D (results for 60 nm and 40 nm clusters are provided in Supplementary Fig. S8). At each point we have optimized the density threshold *minPts* to differentiate from random background noise, similar to the corresponding selection of *minL* in FOCAL3D.

**Figure 4:**
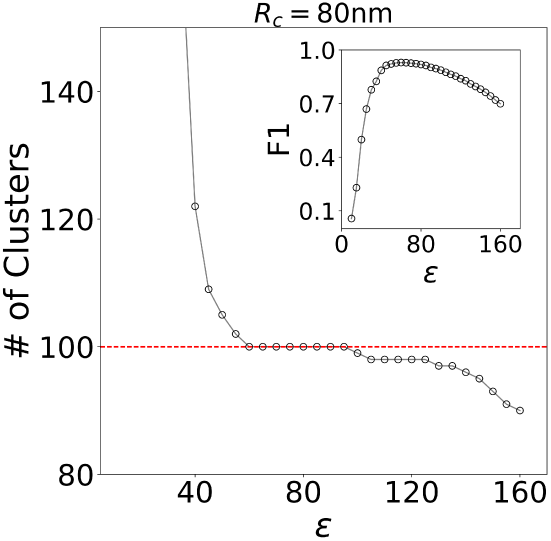
DBSCAN performance at *ζ* = 0.01 for target clusters of radius 80, 60 and 40 nm. Figures display the detected number of clusters vs. the length scale *ϵ* defining the local volume. The dashed line indicates the target number of clusters. Insert: F1 score vs. *ϵ*.

For these simulations, DBSCAN also displays a clear plateau or region of insensitivity to the input parameter defining the local area (i.e., *ϵ*). However, the F1 scores appear to peak at the onset of the plateau, for lower values of *ϵ*, then steeply decline while still within the plateau region. Regardless, the plateau appearing in plots of cluster number vs. *ϵ* indicate a non-biased way to select the appropriate parameters for DBSCAN. As we will show, these regions of insensitivity may disappear in more complex datasets than those we have simulated, leaving the user without a guide for selecting the clustering parameters.

#### 3.2.3 FOCAL3D Performance Under High Noise Conditions

We now consider the performance of FOCAL3D under increasingly noisy conditions, both at *ξ* = 0.05 and *ξ* = 0.20. Our numerical results are displayed in Fig. 5. For large 80 nm clusters and *ξ* = 0.05, signature curves appear in a plot of cluster number vs. *minC*. A steep descent is seen at overly fine grid sizes (Δ = 20 nm), giving way to a plateau near an optimized grid size (Δ* = 40 nm), which then drops below the target for coarser grids (Δ = 90 nm). We find that the region of insensitivity seen at Δ* = 40 nm holds even at very high noise levels (*ξ* = 0.20). FOCAL3D performs quite well for smaller clusters (*R*_*C*_ = 60 nm) under these noisy conditions and appears to actually show less sensitivity to the choice of target grid size. However, the algorithm begins to underestimate the number of target clusters for the smallest clusters we simulated (*R*_*C*_ = 40 nm), resulting in significantly poorer performance as the noise is increased from *ξ* = 0.05 to *ξ* = 0.20.

**Figure 5:**
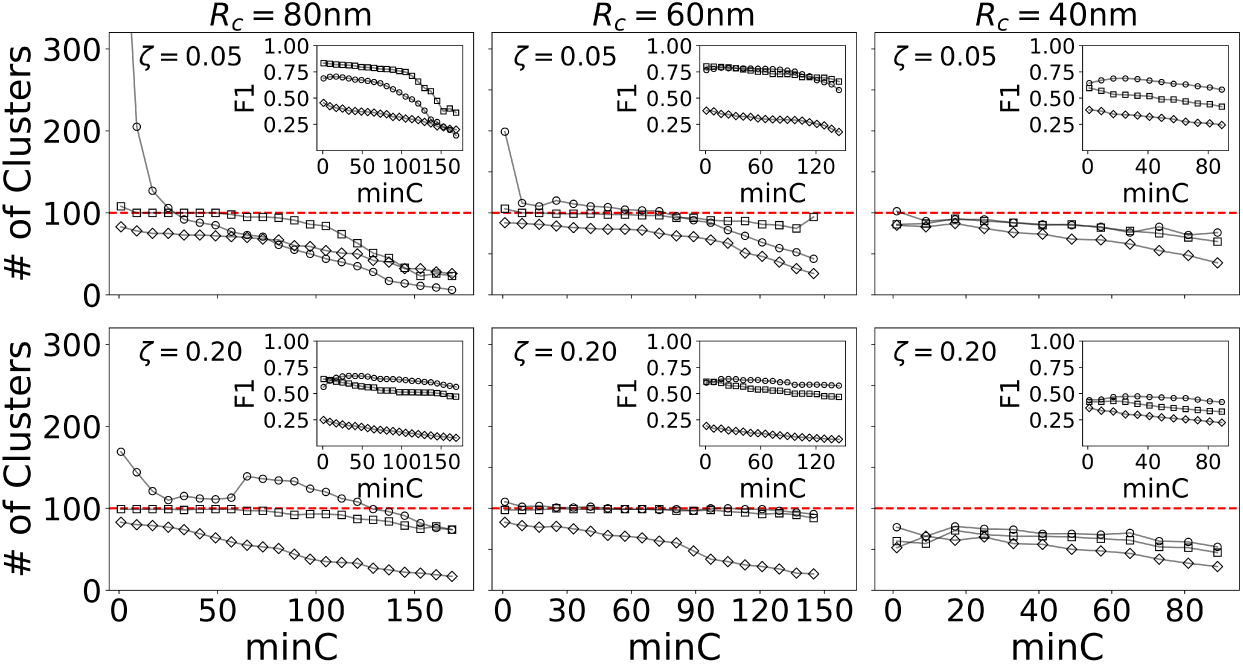
FOCAL3D performance at higher noise levels (*ζ* = 0.05 and *ζ* = 0.20) for 80 nm clusters: Δ = 20 nm (○), Δ = 40 nm (□), Δ = 90 nm (◊); 60 nm clusters: Δ = 20 nm (○), Δ = 30 nm (□), Δ = 80 nm (◊); 40 nm clusters: Δ = 20 nm (○), Δ =40 nm (□), Δ = 60 nm (◊).

Again, at the target grid size Δ*, the F1 scores all display a relatively flat, maximum throughout the corresponding plateau region of *minC*. We note that at these higher noise levels the F1 scores lower, in part, because the background noise in the simulations is randomly distributed throughout the volume. For example, at *ξ* = 0.20, roughly 20% of the points contained within a cluster are considered background noise. Even if the algorithm were to perfectly identify the clusters, it would have no way to discern that these points are noise, resulting in an increased number of false positives.

### 3.3 Clustering of the Nuclear Pore Complex

We now apply our clustering algorithm to 3-dimensional astigmatic dSTORM images of the nuclear pore complex (NPC) ([14, 15, 16]). The images were acquired in wild-type U-2 human osteosarcoma (OS) cells expressing Nup107-SNAP, which was fluorescently labeled with the organic dye Alexa-647 and induced to photoswitch by modifying the imaging buffer. Details on the cell cultures, labeling, fixation, and imaging can be found in ([14]).

We initially partition the dataset focusing on an approximately 2 *µ*m × 2 *µ*m region in the lateral plane centred on the nucleus. Figures 6A-C show the results of the clustering analysis with FOCAL3D. In Figure 6A, we scan through grid sizes ranging from 15-45 nm in steps of 10 nm. The smallest grid size (Δ = 15 nm) shows a steep decline in the number of clusters detected as a function of *minC*. This dependence significantly levels out at a grid size of Δ = 25 nm, with both the extent of the flat region and the estimated number of clusters decreasing for larger choices of the grid size. The discrete, step-like behaviour seen in the curves is simply an artifact of the small number of clusters that we are considering here. We identify Δ* = 25 and choose *minC*\* = 60, which is just at the cusp of the plateau. Figure 6B displays the localizations for this subset of the data in which we can manually identify NPCs by eye. A comparison with the clustering results (Fig. 6C) nicely illustrates the performance of FOCAL3D. Note, the clustering is not strongly dependent upon the choice of grid size for a range of Δ*. For instance, we could have, alternatively, chosen Δ* = 35, which at *minC*\* = 60 identifies roughly 1 less NPC and clusters the localizations in a similar fashion.

**Figure 6:**
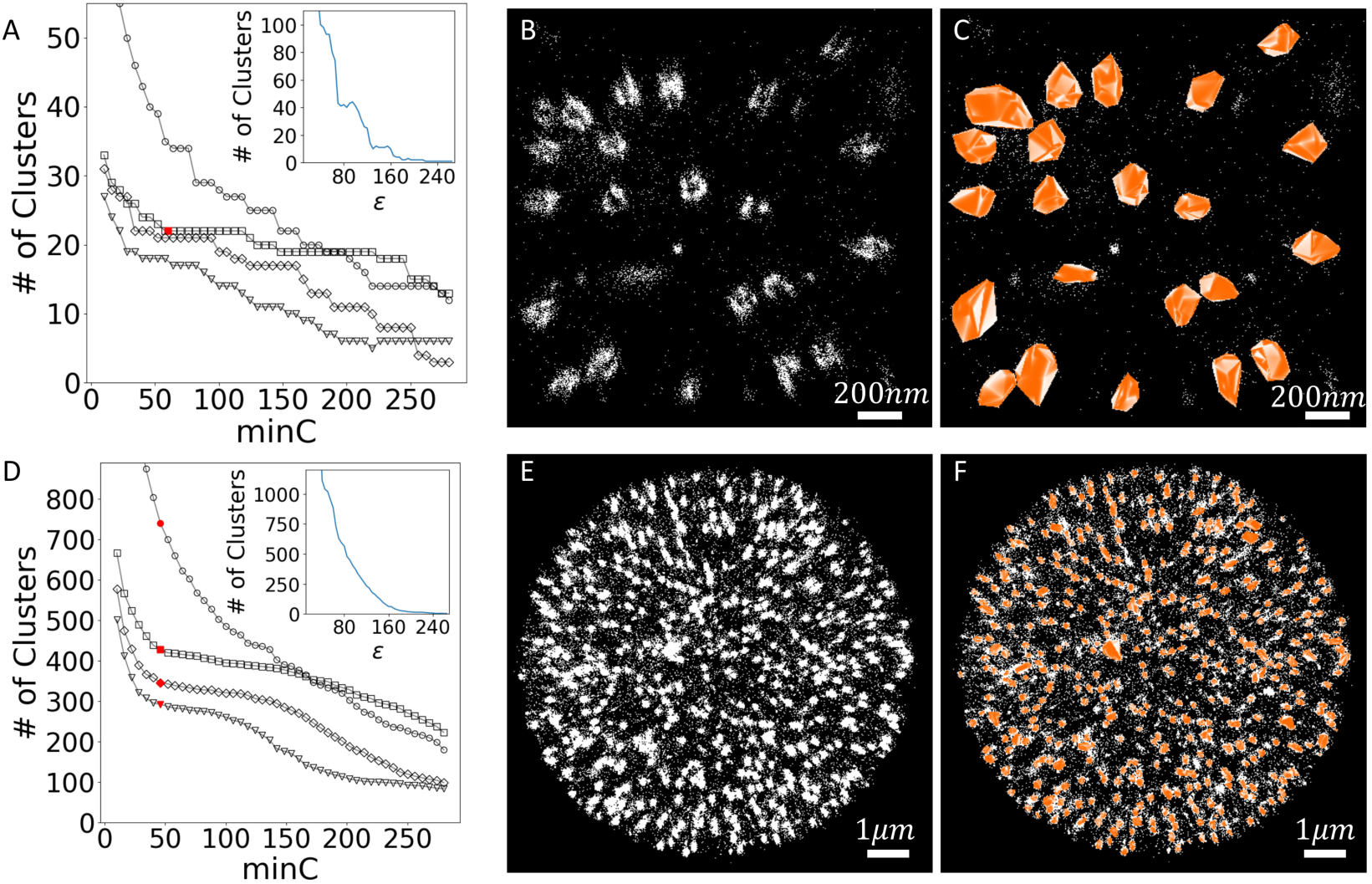
Cluster analysis of NPC dataset in U-2 OS cells. A) FOCAL3D analysis of the number of Clusters vs. *minC*, for the cropped dataset, for grid sizes of Δ = 15 nm (○), 25 nm (□), 35 nm (◊), 45 nm (▽). Insert: Corresponding DBSCAN analysis of the # of Clusters vs. *ϵ*. B) Localization data. C) FOCAL3D clustered dataset at Δ* = 25 nm and *minC*\* = 60. D) FOCAL3D analysis of # of Clusters vs. *minC* for the full dataset. Symbols are the same as in the cropped dataset. Insert: Corresponding DBSCAN analysis of the # of Clusters vs. *ϵ*. E) Localization data. F) FOCAL3D clustering results at *minC*\* = 46 and Δ* = 25.

We also attempted to guide a similar analysis with DBSCAN by generating a plot of the number of detected clusters vs. *ϵ* (insert to Fig. 6A). However, the results showed a steep functional dependence with no clear indication of how to select for *ϵ*. While a number of steps or plateaus appeared in this plot, the most extensive ones occurred far from the target number of clusters (around N=10 and 40).

In Figures 6D-F we consider the full NPC dataset in which a manual enumeration of the NPCs would be difficult. Figure 6D shows an analysis of the detected number of clusters vs. *minC* over the same range of grid sizes (Δ = 15 − 45 nm) as analyzed for the smaller data subset. Once again, an overly fine grid size (Δ = 15 nm) shows a steep dependence on *minC*, with the curves levelling off at Δ = 25 nm, after which the plateau region shrinks and the number of clusters identified for a given *minC* drops. Again, we provide a localization image (Fig. 6E) that can be compared to the clustering results (Fig. 6F). Here we have chosen Δ* = 25 and *minC*\* = 46. A similar analysis for DBSCAN gives no indication of an appropriate way to select for *ϵ* (insert of Fig. 6D).

For this larger dataset, we note that there is a certain level of ambiguity in the choice of Δ*, and the clustering that results is more sensitive to this choice. The effect is primarily due to the close proximity of many of the clusters, a facet that was not explored in our simulations where we explicitly prevented the clusters from overlapping. This can be quantified to some extent by the silhouette score *S*_*C*_ (Supplementary Table S1). For example, in the simulations shown in Fig. 2, optimal clustering resulted in a silhouette score within the range of *S*_*C*_ = 0.80 − 0.85. For the restricted NPC dataset shown in Figs. 6A-B, *S*_*C*_ = 0.53 at Δ* = 25 nm.. However, at the inflection points in Fig. 6D, indicated by the shaded red symbols, we find *S*_*C*_ = 0.48, 0.46, and 0.42 for Δ = 25 nm, 35 nm, and 45 nm, respectively. This indicates that within the image, on average, the predicted clusters are significantly more tightly packed.

FOCAL3D was not designed to distinguish overlapping clusters, so we would expect it to have difficulty with such a dataset. To better understand the behaviour of our algorithm, however, and to further guide us in our parameter selection, we considered the distribution of clusters detected at the cusp of each plateau in Fig. 6D (indicated by the shaded red points in the figure). Starting from the smallest grid size and increasing, the distribution in cluster radii tend to shift toward a peaked distribution, but then develop an increasingly long tail containing larger clusters (Supplementary Fig. S12). The latter effect most likely arises when the algorithm begins to merge the clusters, so the ideal clustering should occur just before this tail develops. This is further justification that our choice of Δ* = 25 was appropriate. As a final check, the peak of the distribution in cluster sizes agrees with a Ripley’s K-function analysis (Supplementary Fig. S13 and Fig. S14).

## 4 Discussion

As SMLM continues to push the resolution limit obtainable by light microscopy, efficient algorithms need to be developed that can cope with the concomitantly larger datasets that will be generated. In this manuscript, we have presented an extension of our 2-dimensional Fast Optimized Clustering Algorithm for Localization Microscopy (FOCAL) ([13]) to 3-dimensions. FOCAL3D is a density based algorithm, so directly identifies clusters in noisy SMLM images, and shows significant performance improvements when compared to the classical density based algorithm DBSCAN (scaling like *n*_*l*_ compared to *n*_*l*_ log *n*_*l*_). Moreover, unlike DBSCAN, the algorithmic parameters that determine FOCAL3D’s performance can be systematically tuned such that, within a constrained range of parameters, the clustering is only weakly dependent upon the exact choice of these parameters.

While the speed gains of working on a grid can be considerable, for small clusters that necessitate an excessively fine grid, the computational cost of FOCAL3D can outweigh that of DBSCAN. As we saw in our simulations, the region of parametric insensitivity is reduced making it harder to optimize the cluster detection. Likewise, FOCAL3D may have issues analyzing dense SMLM image reconstructions, particularly when the clusters begin to overlap. Future extensions to FOCAL3D may alleviate these issues, such as incorporating a segmentation algorithm to differentiate anomalously large, dense clusters of localizations.

FOCAL3D is designed to automate and rapidly perform an analysis of large 3-dimensional SMLM data sets. Likewise, the resulting clustering may be used as a way to filter out noise in SMLM image reconstructions, retaining only the features of interest. FOCAL3D should serve as a useful addition to the set of quantitative techniques now available for super-resolved microscopy, providing a foundation for further analysis of intracellular organization, protein assemblages, and spatial patterning.

## Acknowledgements

We thank Dr. Jonas Reis at EMBL in Heidelberg, Germany for sharing 3D SMLM data on the nuclear pore complex. We also thank Muhammad Kamal for developing a graphical user interface for FOCAL3D. Andreas Hilfinger and Emiel Visser provided valuable feedback on the manuscript.

## Supporting Information

### 1 Clustering Optimization

#### 1.1 Flowchart of FOCAL3D optimization

**Figure S1:**
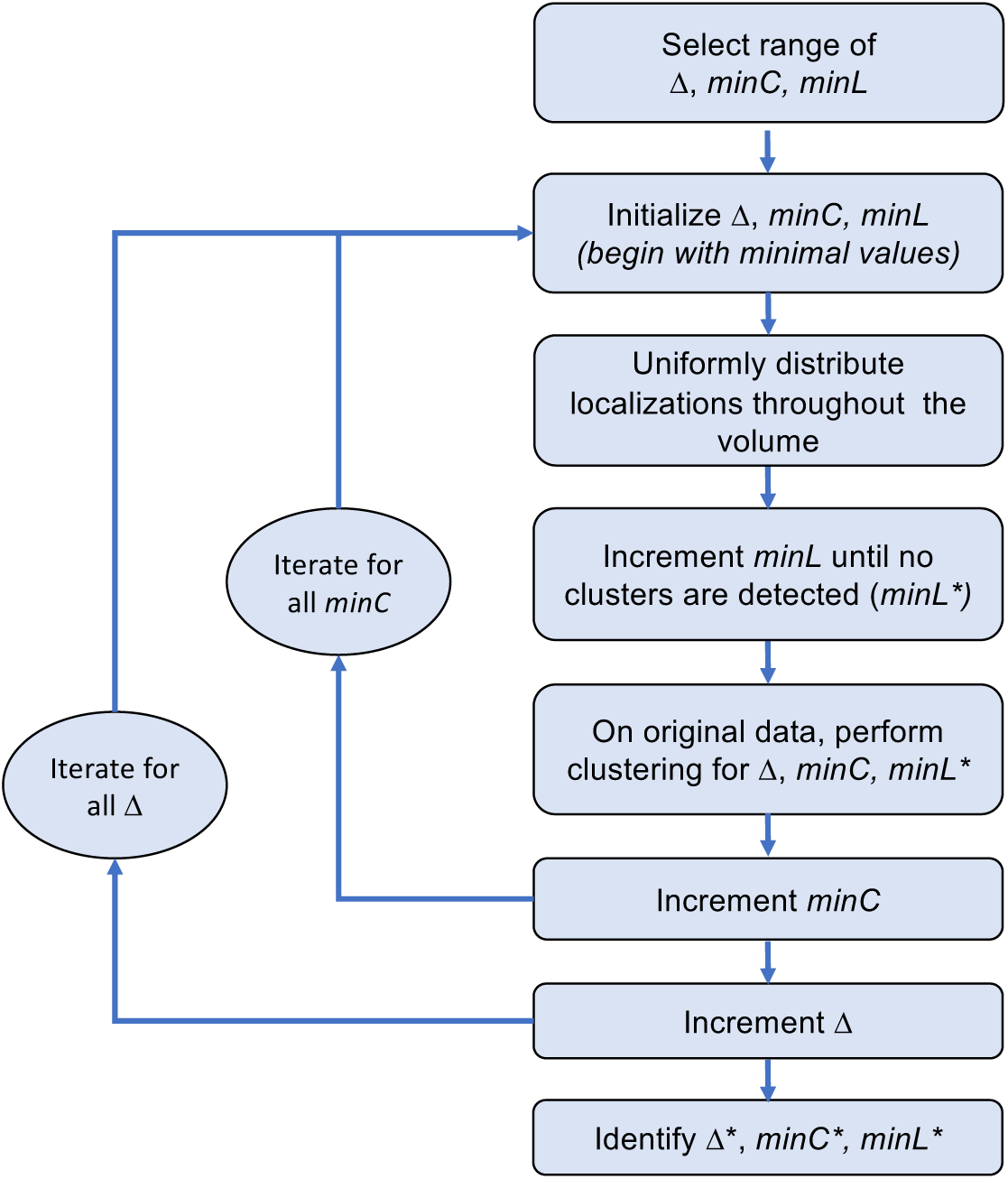
A flowchart illustrating the discussion in Section 2.2 on optimizing the parameters in FOCAL3D.

#### 1.2 Lower bound of *minL* in FOCAL3D

To save computation time in optimizing *minL*, we can narrow the search range by estimating a lower bound. We find an optimal *minL* by randomly scattering all *N*_*total*_ localizations within the image volume *V*, so that the density of localizations is *N*_*total*_*/V*. For a given grid size Δ, the volume occupied by a voxel is *δV* = Δ^3^ and the average number of localizations per voxel is 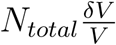. Since the image is initially enhanced by locally summing all neighbouring voxels, we should multiply by the voxel connectivity (3^3^ − 1 = 26 in 3D or 3^2^ − 1 = 8 in 2D). So, for the 3D case, we find a lower bound of:

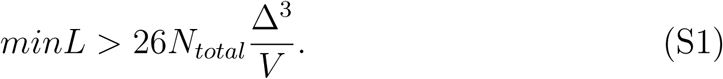

**Figure S2:**
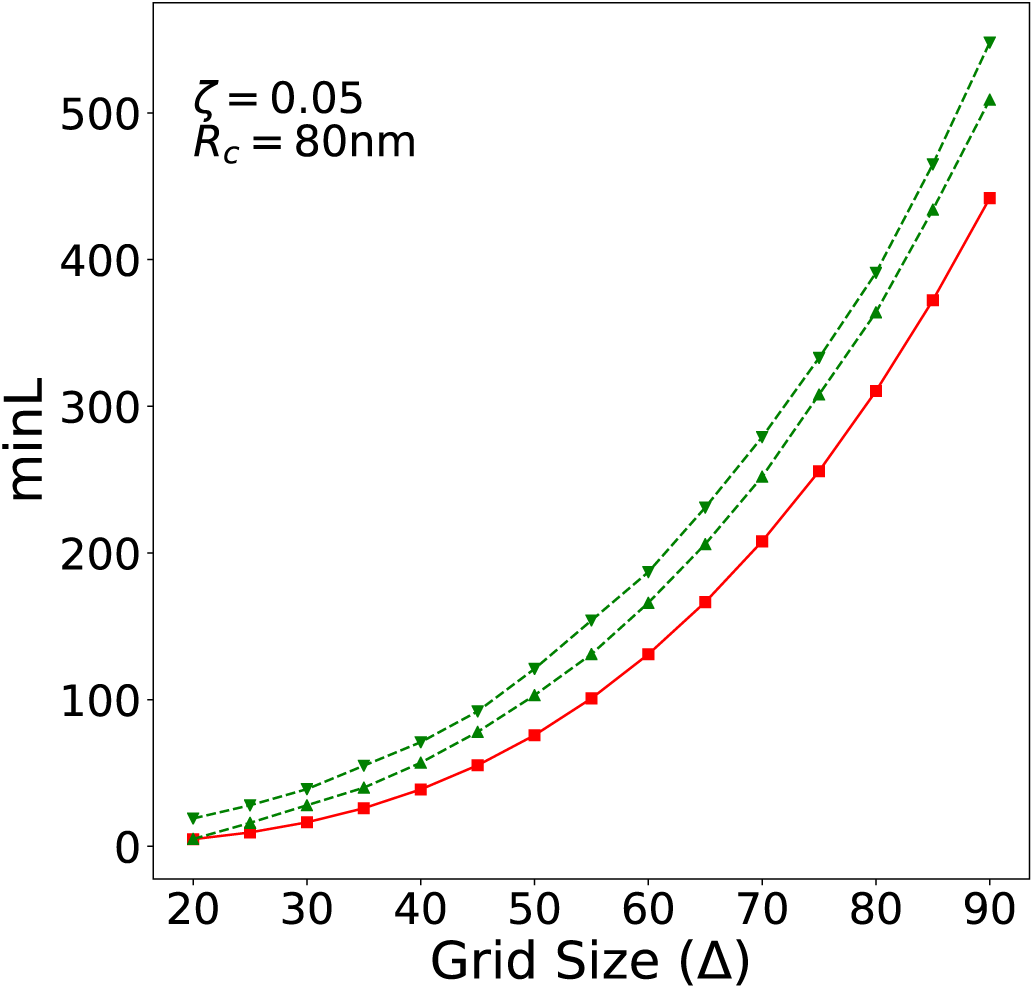
Estimating the lower bound of *minL* in FOCAL3D. *ζ* = 0.05 and a cluster size of 80±16 nm within a 5 *µ*m×5 *µ*m×5 *µ*m simulation volume. Estimated lower bound (□), *minL*\* at *minC* = 5 (▽):, *minL*\* at *minC* = 141 (Δ)

#### 1.3 Lower bound of *minPts* in DBSCAN

We can also set a lower bound for *minPts* in DBSCAN. Similar to the case for *minL*, we can estimate the average number of randomly scattered localizations we expect in a sphere of radius *ϵ*:

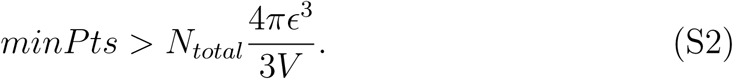

**Figure S3:**
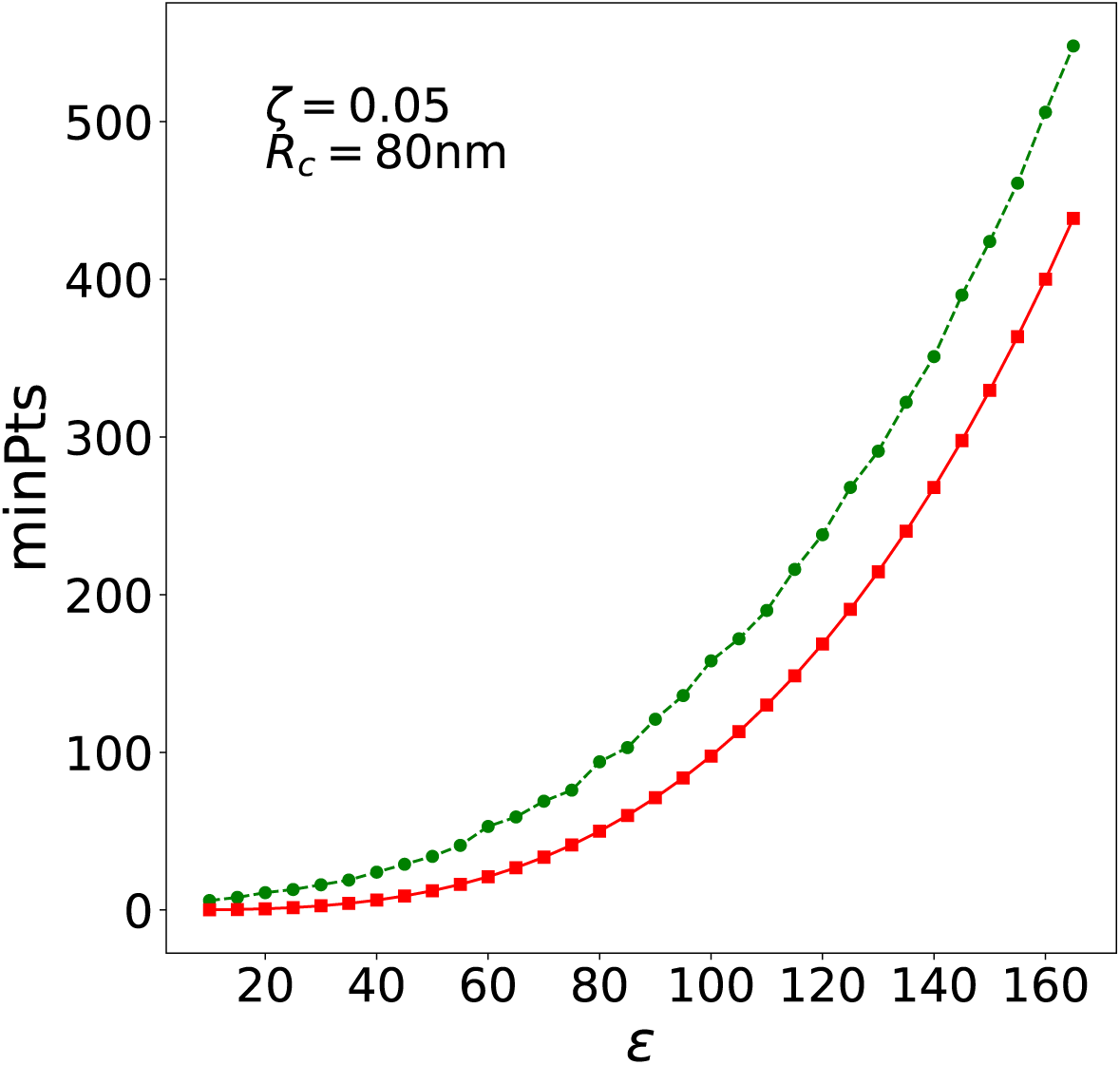
Estimating the lower bound of *minPts* in DBSCAN. *ζ* = 0.2 and a cluster size of 80 ± 16 nm within a 5 *µ*m×5 *µ*m×5 µm simulation volume. Estimated lower bound (□), *minPts*\* (•).

### 2 Simulation Details

#### 2.1 Distribution of Number of Localizations per Fluorophore

Experimentally, fluorophores typically used in single-molecule localization microscopy have an average ON time which follows an exponential distribution:

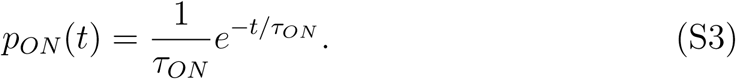

If we work in units of the exposure time *τ*_*exp*_, and identify *n* ≈ *t*/*τ*_*exp*_ as the number of localizations that will be detected during a single emission period due to temporal binning by the camera, we can rewrite Eq. S3 as:

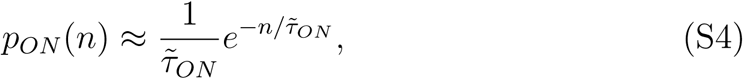

where 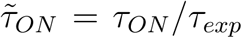. Moreover, almost all photoswitchable fluorophores blink, where we define a blink as a reactivation of the emissive state of the fluorophore. The number of blinks is commonly observed to follow an geometric distribution

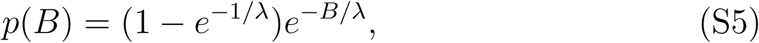

where *λ* is the characteristic number of blinks.

We are interested in the distribution of total localizations *n* from a single fluorophore. Let’s first consider the distribution of occurrences of

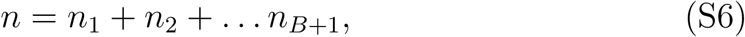

where *n*_1_, *n*_2_,…*n*_*B*+1_ are the number of localizations observed for a given number of blinks *B*. Since the sum of *B*+1 independent identical exponential variables is a Gamma function (i.e., 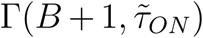), from Eq. S4 and Eq. S5, we can derive the probability distribution for the number of localizations expected from a fluorophore that blinks *B* times:

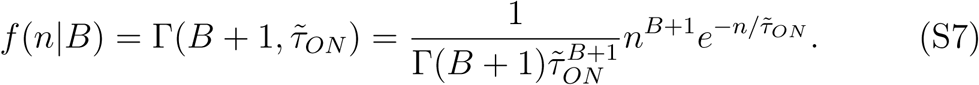

Finally, we weigh the probability distribution *f*(*n*|*B*) by the probability of observing B blinks to obtain the distribution of the number of localizations per fluorophore *p*(*n*):

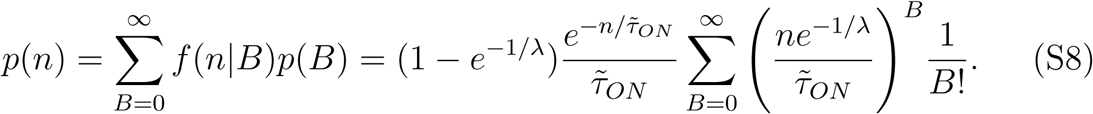

The sum is just the Taylor series expansion of *e*^*x*^ where 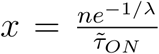 in this case. Thus, the distribution of the number of localizations per fluorophore is

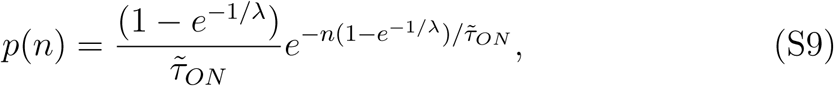

which is an exponential distribution with a characteristic number of localizations per fluorophore of 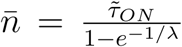. Since localizations are a discrete quantity we can discretize the exponential distribution into a geometric distribution as

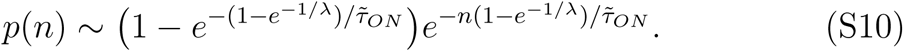

#### 2.2 Distribution of Localization Precision

We can model the distribution of localization precisions by taking into account the distribution of the total number of photons per blink:

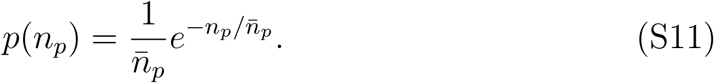

An approximate lower bound for the localization precision is given by 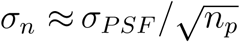. From this we can find a model distribution of the localization precision:

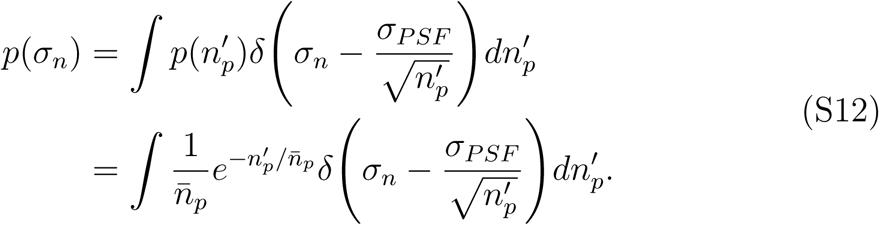

With the substitution 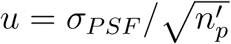, this gives:

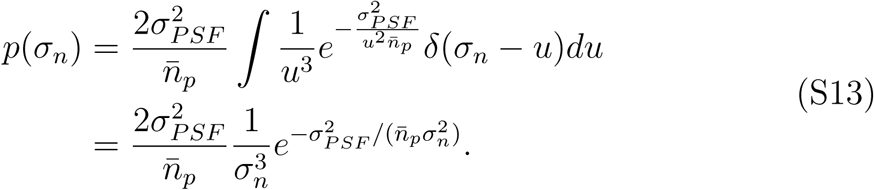

The mode of this distribution can be calculated from the maximum likelihood estimate as:

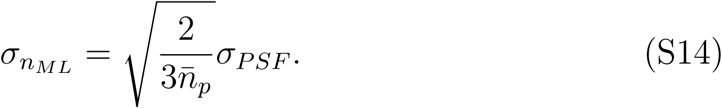

The mean of this distribution is:

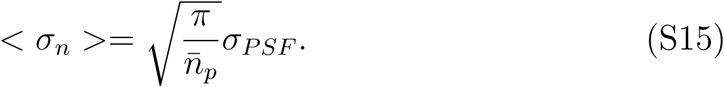

These relations are useful in scaling the axial localization precision with respect to the lateral precision. For example, by taking double the PSF width in the axial direction as in the lateral dimensions, we effectively shift the peak and mean of the distribution by a factor of 2.

**Figure S4:**
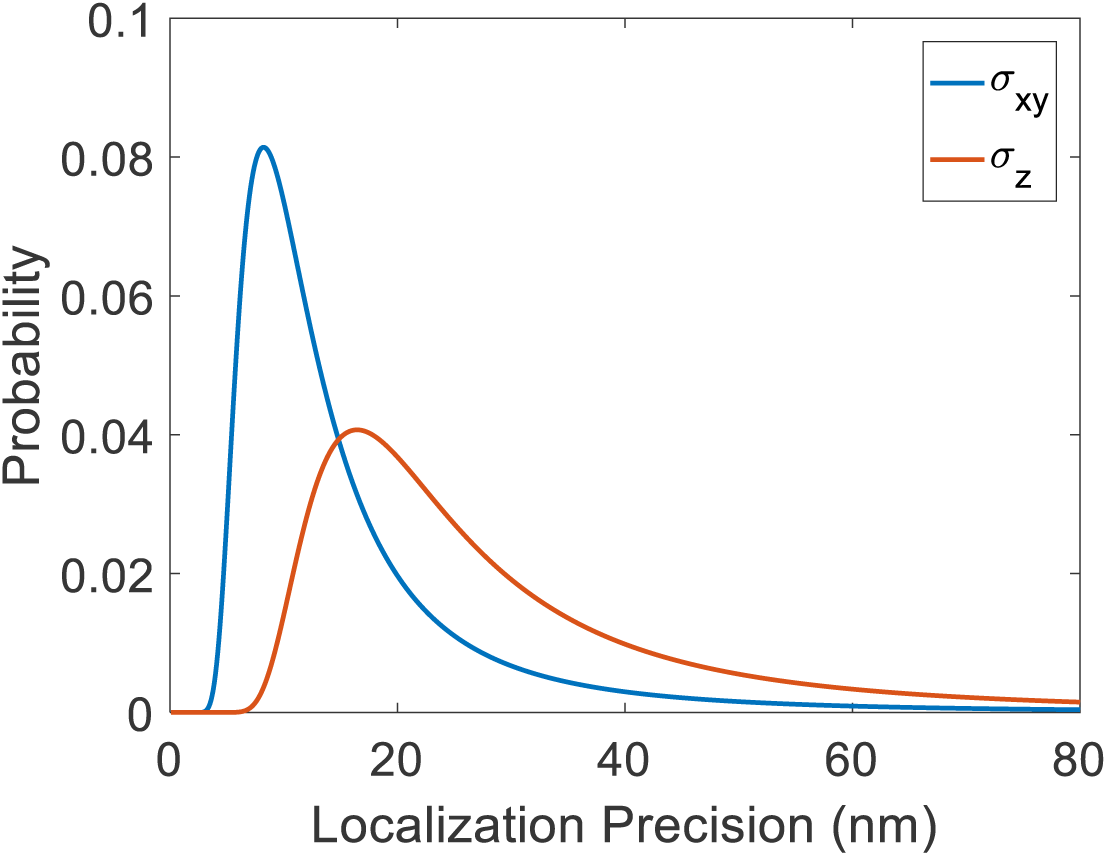
Theoretical localization precision distributions with 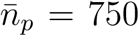, and a point-spread function width of 390 nm in the lateral direction and 780 nm in the axial direction

To use this for simulations, we use inverse transform sampling. The cumulative distribution function (CDF) of the localization precision is:

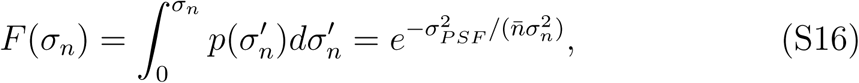

and the inverse CDF is:

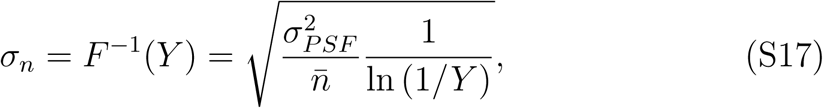

so if *Y* ∼ *U*(0, 1), then *σ*_*n*_ *∼ p*(*σ*_*n*_).

### 3 Ground Truth Cluster Sizes

**Figure S5:**
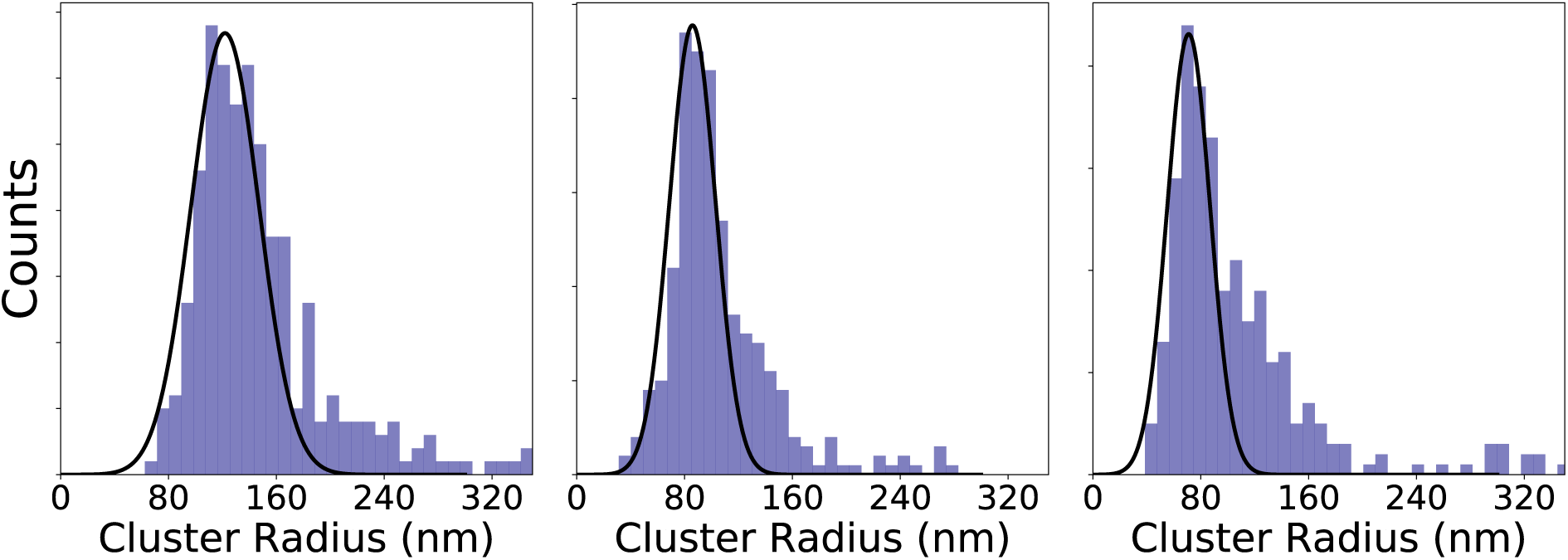
Simulated ground truth cluster radii obtained by convex hull for A) 80 ± 16 nm clusters, B) 60 ± 12 nm clusters, C) 40 ± 8nm clusters. Solid black line is a Gaussian fit to the peak of the distribution.

### 4 Insensitivity to Grid Selection

There is some flexibility on the choice of optimal grid size as illustrated in these graphs, which correspond to the results from Fig. 2 of the main text.

**Figure S6:**
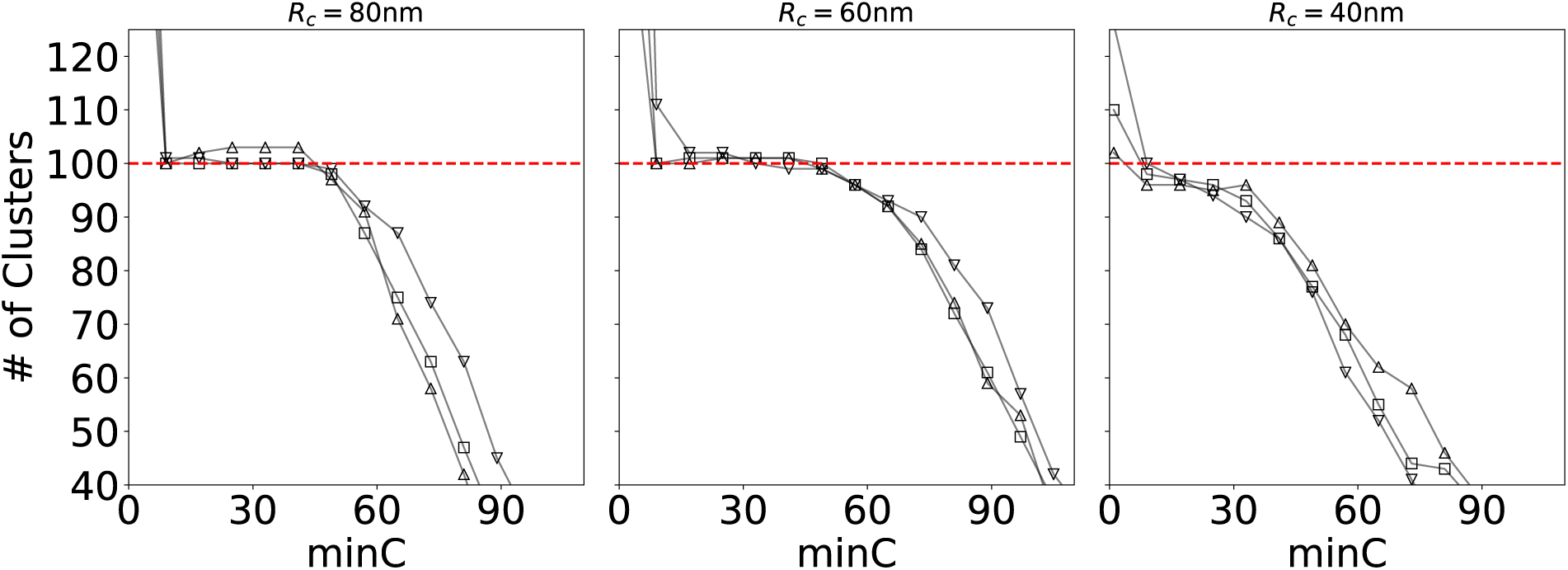
FOCAL3D performance for simulations at a noise level of *ζ* = 0.01. For *R*_*c*_ = 80 nm: Δ = 35 nm (▽), Δ = 40 nm (□), Δ = 45 nm (▵). For *R*_c_ = 60 nm: Δ = 25 nm (▽), Δ = 30 nm (□), Δ = 35 nm (▵). For *R*_c_ = 40 nm: Δ = 30 nm (▽), Δ = 40 nm (□), Δ = 50 nm (▵)

### 5 Run-Time Scaling

We can compare the time performance of FOCAL3D and DBSCAN as a function of their respective parameters or as a function of the number of localizations (*n*_*l*_) in the data set. To perform a fair comparison, we match the local volumes employed by the two algorithms by the relation *minC* * Δ^3^ = (4*π*/3)*ϵ*^3^.

**Figure S7:**
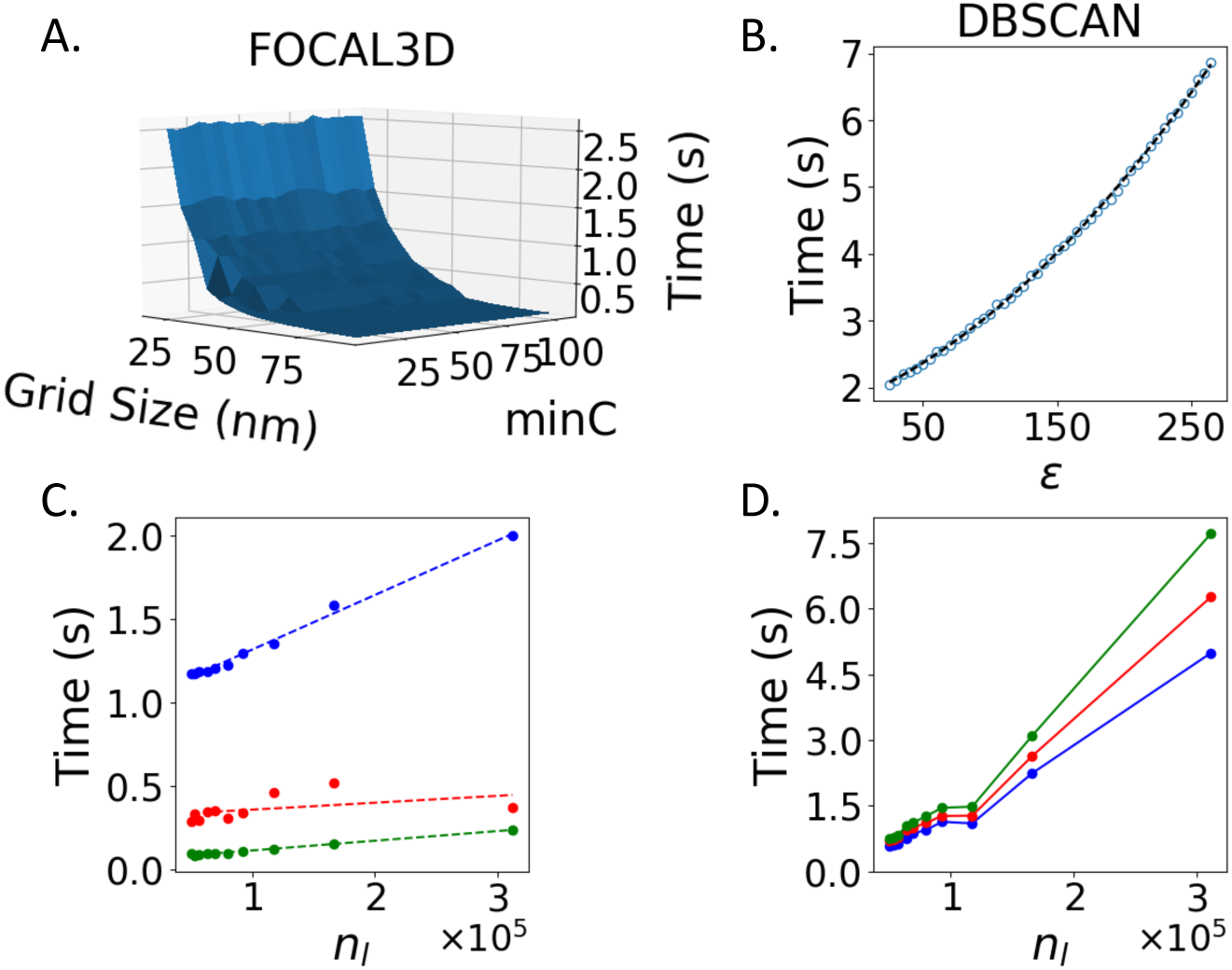
A) Time dependence of FOCAL3D for simulated data of 80 nm clusters (*ζ* = 0.01, *n*_*l*_ = 166274 localizations) as a function of grid size Δ and *minC*. The time quickly decays for increasing Δ, but remains essentially constant as a function of *minC* for a fixed grid size. B) Time dependence of DBSCAN for simulated data of 80 nm clusters (*ζ* = 0.01, *n*_*l*_ = 166274 localizations) as a function of *ϵ*. The run-time quadratically increases for increasing *ϵ* (dashed curve). C) Time dependence of FOCAL3D for a varying number of localizations at fixed *minC* = 15: Δ = 25 nm (blue), Δ = 45 nm (red), Δ = 65 nm (green). Dashed lines correspond to linear fits. D) Time dependence of DBSCAN for varying number of localizations at different *ϵ* corresponding, roughly, to the local volume of FOCAL3D: *ϵ* = 40 nm (blue), *ϵ* = 70 nm (red), *ϵ* = 100 nm (green).

### 6 DBSCAN performance for smaller cluster sizes (*ζ* = 0.01)

**Figure S8:**
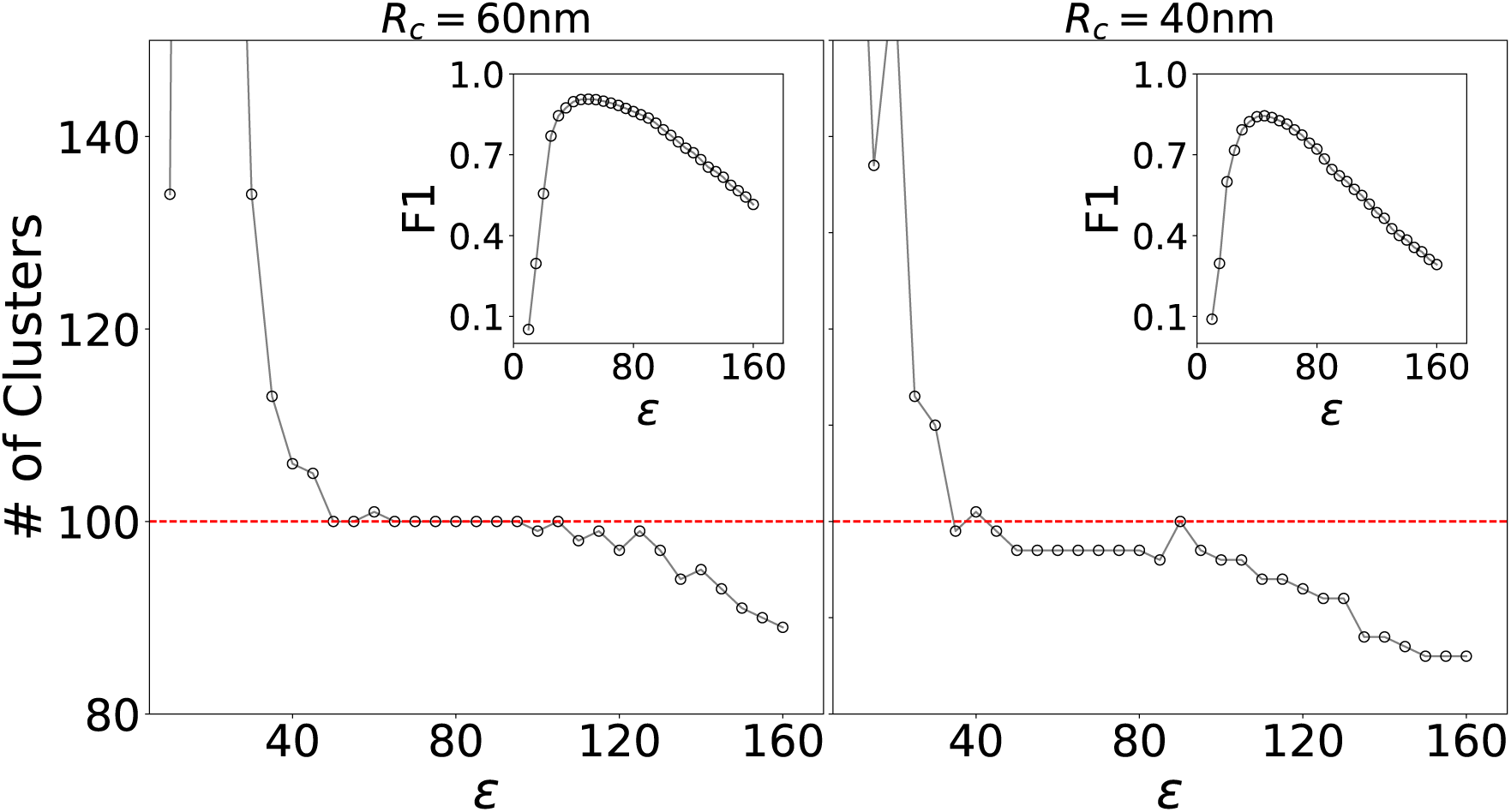
DBSCAN performance at lower grid sizes (60 nm and 40 nm) and moderate noise *ζ* = 0.01

### 7 Precision and Recall

#### 7.1 FOCAL3D (*ζ* = 0.01 Simulations)

**Figure S9:**
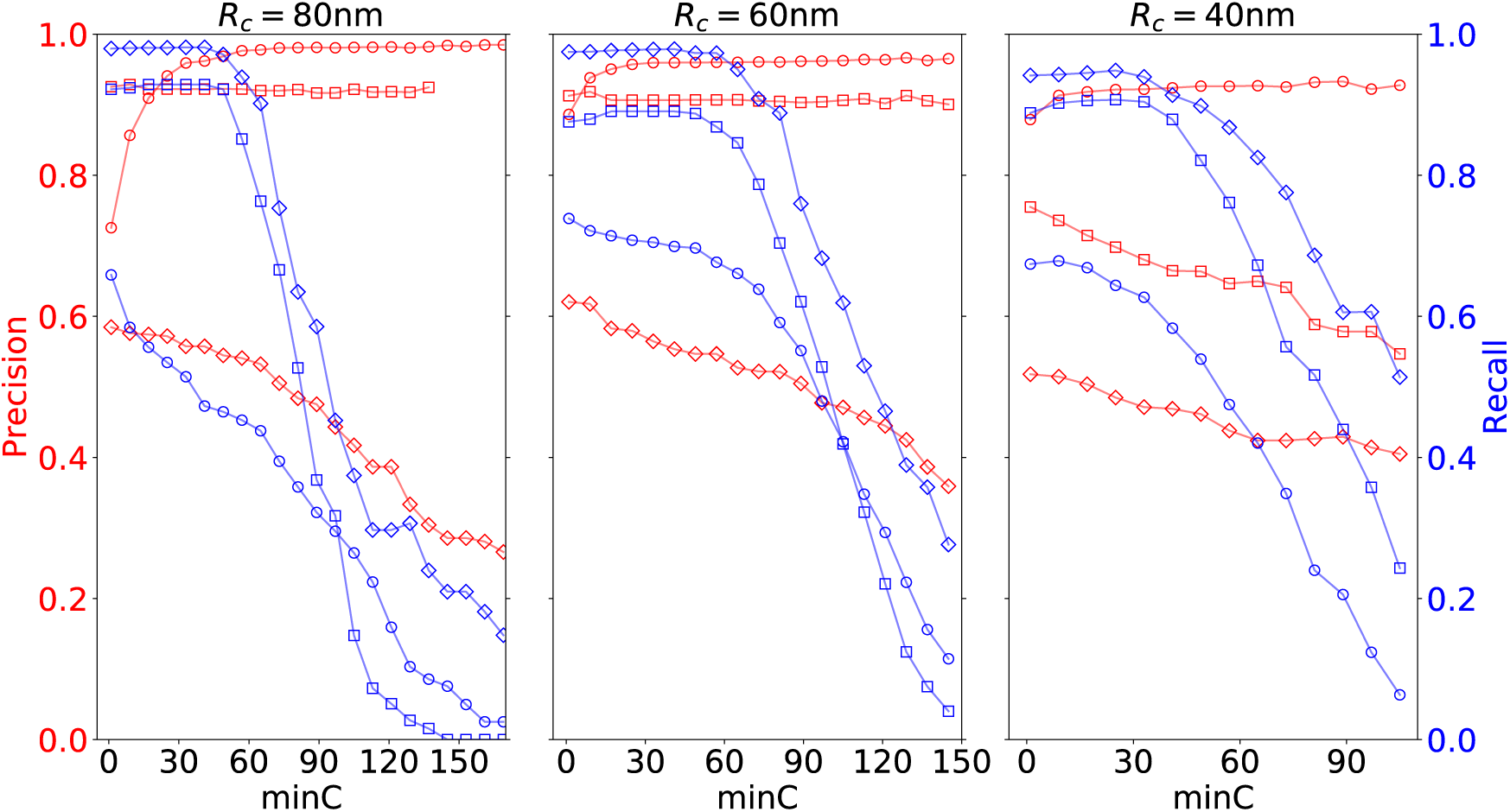
Precision and Recall scores used to calculate the F1 scores in the insets of Figure 3 of the main text

#### 7.2 DBSCAN (*ζ* = 0.01 Simulations)

**Figure S10:**
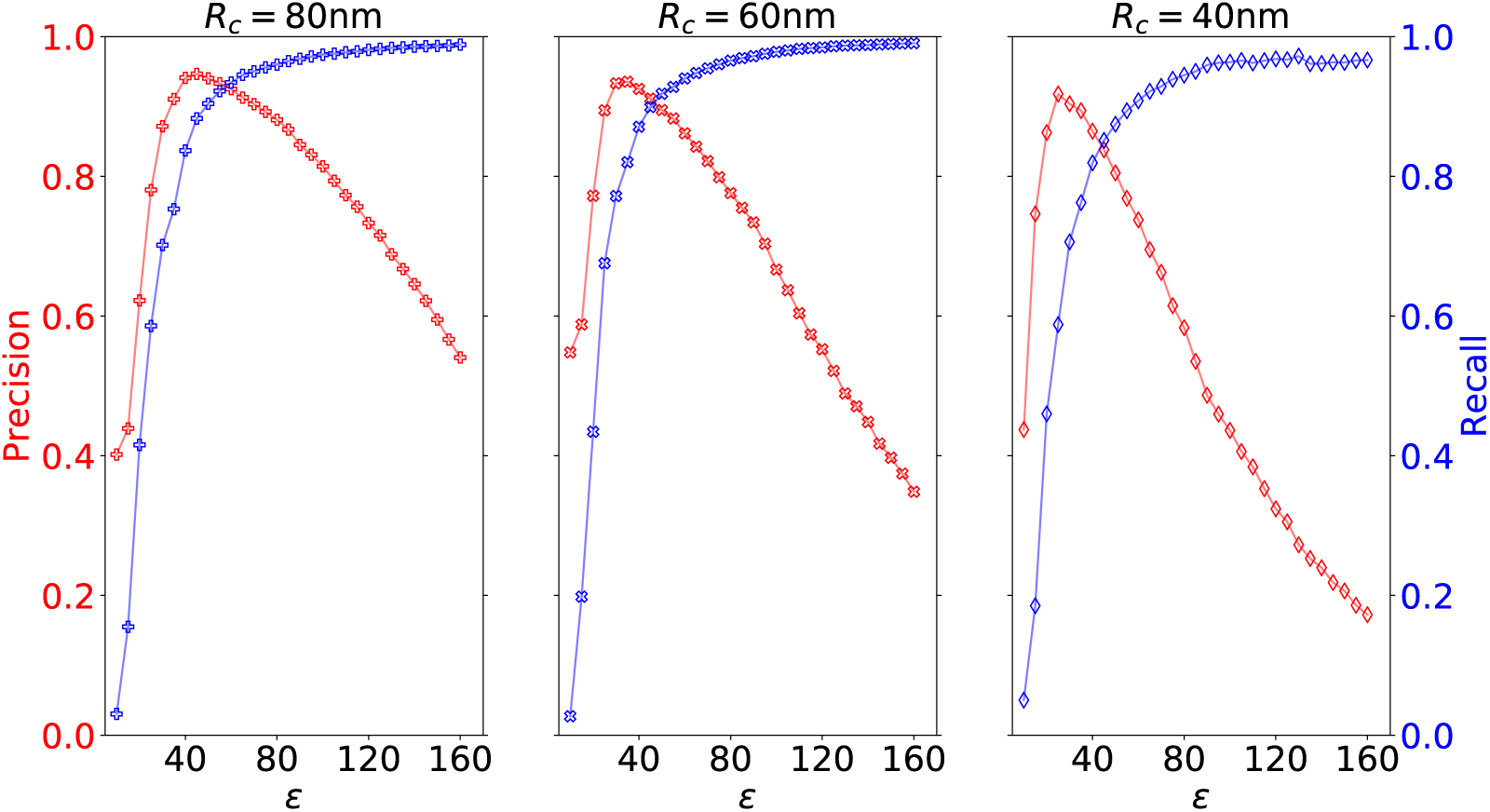
Precision and Recall scores used to calculate the F1 scores in the insets of Figure 4 of the main text

#### 7.3 FOCAL3D (*ζ* = 0.05 and *ζ* = 0.20 Simulations)

**Figure S11:**
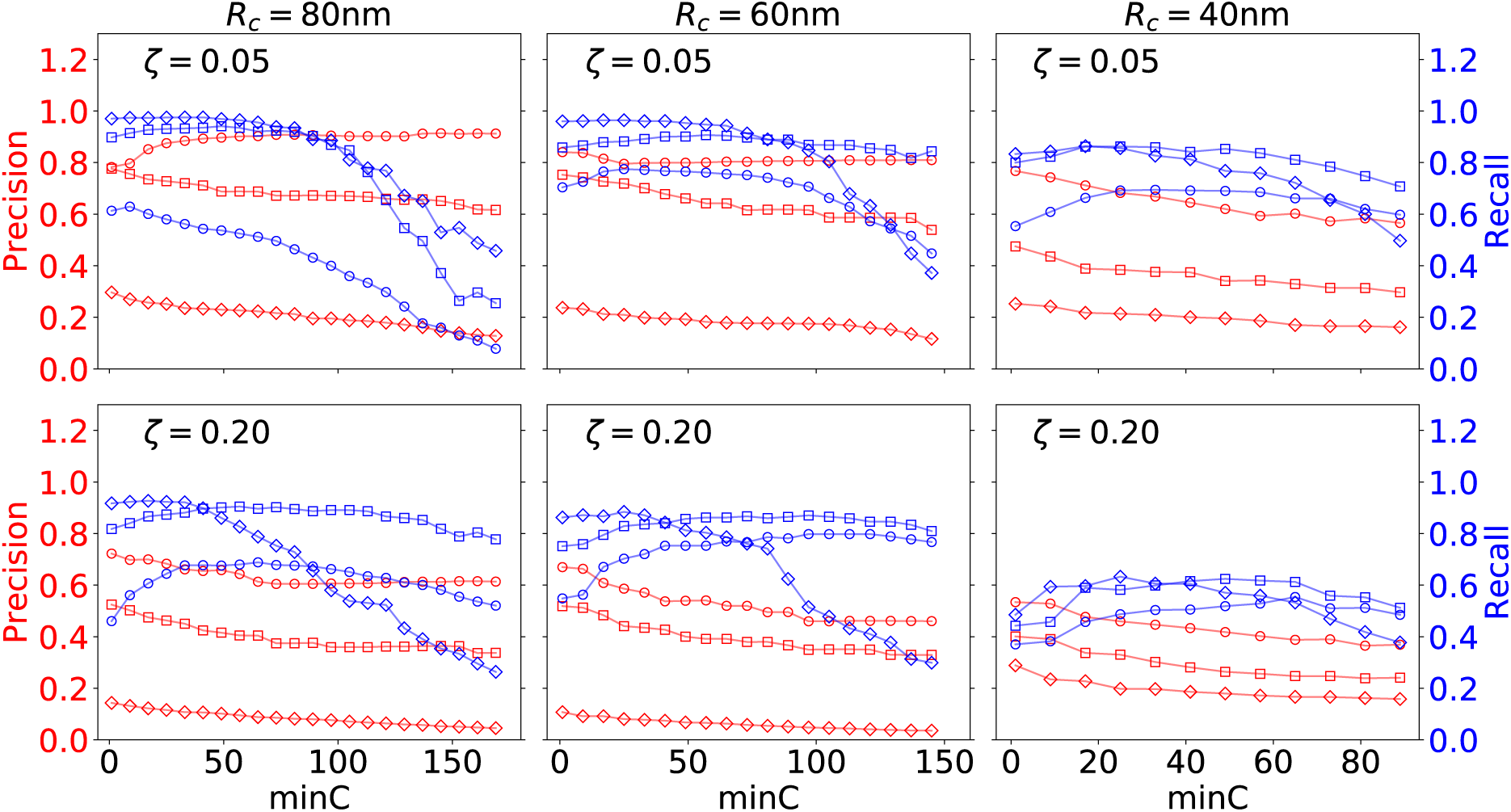
Precision and Recall curves used to calculate the F1 Score for *ζ* = 0.05 and *ζ* = 0.20. Shown are the results for cluster sizes of 80 nm, 60 nm and 40 nm in Figure 5 of the main text.

### 8 Nuclear Pore Complex Analysis

#### 8.1 Cluster radius vs. grid spacing

**Figure S12:**
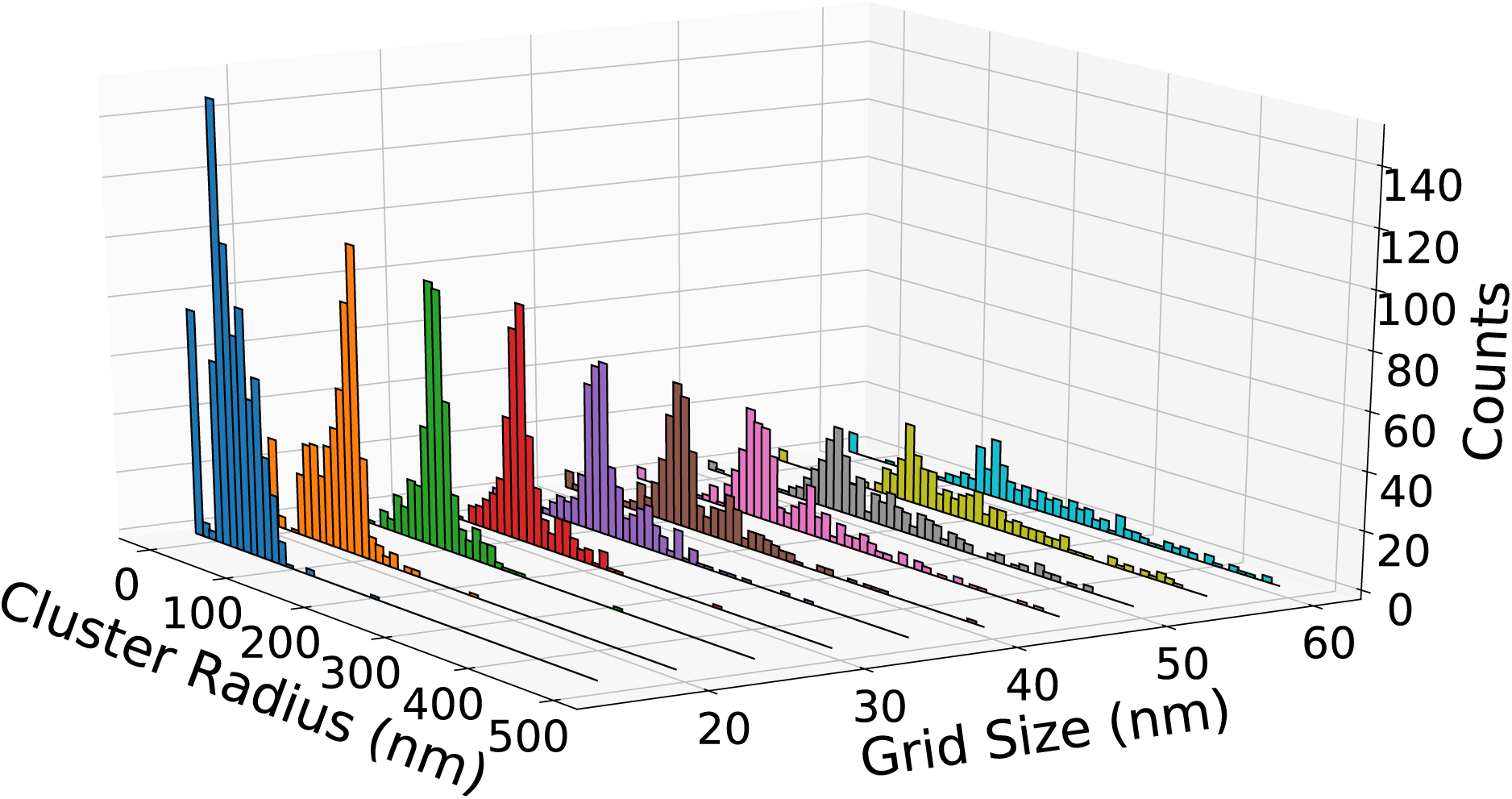
Effective cluster radii for NPC dataset at different grid sizes (evaluated at *minC* = 46 in Fig. 6). For increasing grid size, the distribution in cluster radii first shifts toward a peaked distribution. Then for larger grids, this peaked distribution gradually diminishes while increasingly extending a long tail (indicating large clusters). This is due to separate, but neighbouring, NPCs being grouped into the same cluster

#### 8.2 Cluster radius vs. *minC*

**Figure S13:**
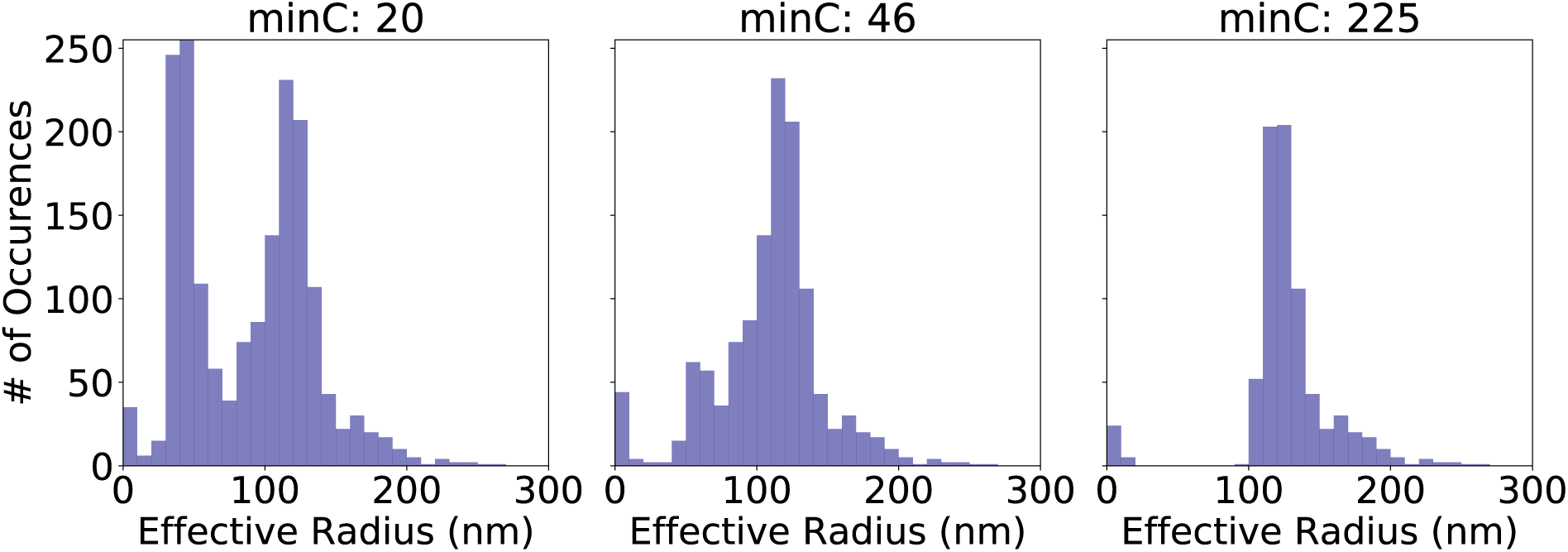
Effective cluster radius, determined by convex hull, for NPC dataset at different values of minC (Δ* = 25 nm). A similar behaviour is observed as in the simulations. For small *minC*, several false small clusters are identified. About some optimal value, we find a single peaked distribution. And for too large a choice of *minC*, reasonable, smaller clusters begin to get cut by the size threshold. The average cluster size for a *minC* of 46 is 109 ± 39

#### 8.3 Effective cluster radius from Ripley’s H-function

**Figure S14:**
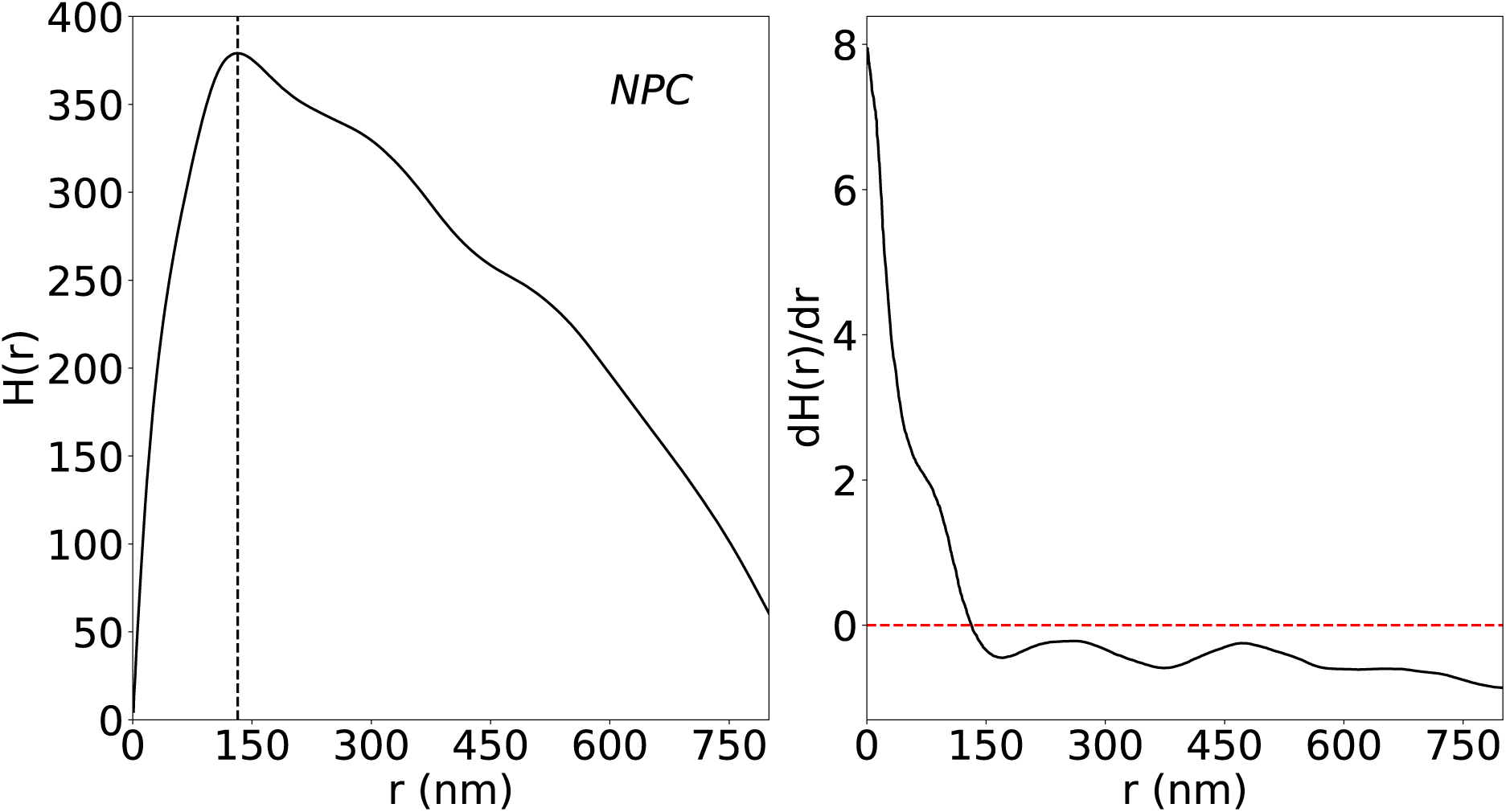
Ripley’s H-function (left) and its derivative (right) for the NPC data set. The cluster size is estimated to be about 132 nm.

#### 8.4 Silhouette scores

**Table 1:**
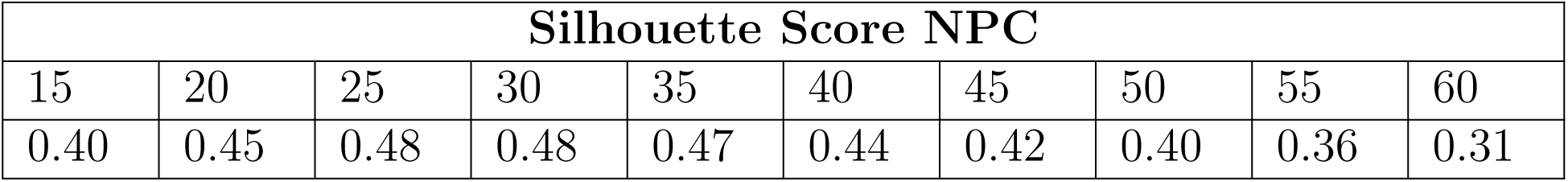
Silhouette scores of the clusters found by FOCAL3D in the NPC data set. There is a maximum at a grid size of around 25-30 nm. The score decreases at high grid sizes as neighbouring NPCs increasingly merge.

| Silhouette Score NPC |  |  |  |  |  |  |  |  |  |
| --- | --- | --- | --- | --- | --- | --- | --- | --- | --- |
| 15 | 20 | 25 | 30 | 35 | 40 | 45 | 50 | 55 | 60 |
| 0.40 | 0.45 | 0.48 | 0.48 | 0.47 | 0.44 | 0.42 | 0.40 | 0.36 | 0.31 |

